# Spatial kinetics and immune control of murine cytomegalovirus infection in the salivary glands

**DOI:** 10.1101/2024.02.22.581694

**Authors:** Catherine Byrne, Ana Citlali Márquez, Bing Cai, Daniel Coombs, Soren Gantt

## Abstract

Human cytomegalovirus (HCMV) is the most common congenital infection. Several HCMV vaccines are in development, but none have yet been approved. An understanding of the kinetics of CMV replication and transmission may inform the rational design of vaccines to prevent this infection. The salivary glands (SG) are an important site of sustained CMV replication following primary infection and during viral reactivation from latency. As such, the strength of the immune response in the SG likely influences viral dissemination within and between hosts. To study the relationship between the immune response and viral replication in the SG, and viral dissemination from the SG to other tissues, mice were infected with low doses of murine CMV (MCMV). Following intra-SG inoculation, we characterized the viral and immunological dynamics in the SG, blood, and spleen, and identified organ-specific immune correlates of protection. Using these data, we constructed compartmental mathematical models of MCMV infection. Model fitting to data and analysis indicate the importance of cellular immune responses in different organs and point to a threshold of infection within the SG necessary for the establishment and spread of infection.

**Author Summary:** Cytomegalovirus (CMV) is the most common congenital infection and causes an enormous burden of childhood disease. To gain insight into the immune requirements for controlling infection, we used a mouse model to reproduce characteristics of natural CMV infection, employing a low viral inoculum, and delivering the virus to the salivary glands (SG), a key site of CMV replication. Our results provide detailed data on the spatial and temporal spread of infection throughout the body and identify key immune correlates of the control of viral replication. By translating these findings into mechanistic mathematical models, we revealed the importance of organ-specific immune responses, particularly the requirement of TNF-*α* and IFN-*γ* to control infection within the salivary glands. Furthermore, our mathematical modeling allowed us to compare known characteristics of human CMV infection related to infection establishment and spread to those predicted in mice, underscoring the suitability of the MCMV model to study its human homologue. These insights provide guidance for developing targeted vaccines to prevent CMV infection and disease.

## Introduction

Human cytomegalovirus (HCMV) is a *β* herpesvirus that infects the majority of the world’s population (1). HCMV establishes life-long infection, primarily acquired via mucosal exposure to virus shed in body fluids, such as saliva, urine, and breast milk, of infected individuals (2,3). HCMV is also the most common congenital infection, occurring in roughly 0.5% of all live births in high income countries, and even more frequently in low and middle-income countries (4). A major driver of congenital infection is transmission from young children, who persistently shed virus at high levels after acquiring HCMV infection, to pregnant women (5,6). While a tremendous amount of research has been dedicated to HCMV vaccine development, clinical trials of candidates performed to date have demonstrated, at most, around 50% protection against HCMV acquisition and have not been approved for use (7–11). However, a recent study by our group indicates that even modestly protective vaccines may be highly effective at decreasing congenital infection if given to young children, due to their ability to reduce viral shedding and transmission to pregnant women (12). As such, a better understanding of the determinants of the intensity and duration of viral shedding would be valuable to inform the development of vaccines to prevent HCMV transmission.

The murine (M)CMV model facilitates studies of these viral dynamics and immune control (13–20). MCMV and HCMV genomes share a high degree of sequence homology and MCMV infection recapitulates many features of its human counterpart (21,22). However, most MCMV experiments have involved inoculating mice with high doses of virus via the intraperitoneal (IP) or intravenous (IV) route of administration (ROA) to ensure infection, rather than simulating the typical conditions of a natural CMV infection involving mucosal exposures to lower quantities of virus (13,14,23).

HCMV infection is most often acquired orally, and viral replication in the salivary glands (SG) is detected early in HCMV infection (24). Thus, low-dose MCMV inoculation of the SG may have particular relevance for natural HCMV exposure. HCMV shedding in saliva tends to occur at higher levels and is more prolonged than in other anatomic sites during primary infection and reactivation from latency (25–27). In mice, the SG also appear to represent a distinct compartment of infection in which active MCMV replication lasts weeks longer than in other tissues (13,28,29). Studies have shown that MCMV effectively prevents major histocompatibility (MHC) class I expression on infected SG cells, thus abrogating recognition and destruction by CD8 T cells, which helps to explain persistent, high-level viral shedding in saliva (30). Rather, CD4 T cells eventually control infection in the SG through the production of the cytokines interferon (IFN)-*γ* and tumour necrosis factor (TNF)-*α*, which inhibit viral replication (20,30–32).

Different immune responses in the SG compared to the rest of the body may also explain why MCMV inoculations to this site have been shown to disseminate less frequently to the rest of the body, compared to the IP or intranasal (IN) ROA (13,14). Indeed, human cohort studies by our group also suggest that oral HCMV replication is often self-limiting, and dies out before systemic dissemination and establishment of latent infection can occur, leading to a low within-host reproductive number (*R*_0_) (24,33). Neither the within-host *R*_0_ of MCMV nor the determinants of viral persistence in, or spread from, the SG have been defined.

To address the requirements for establishing infection, immune control at different anatomic sites, and spread from the SG, we performed low-dose MCMV intra-(I)SG infection experiments, collecting high-resolution spatial and temporal data on viral spread and immune response. With these data, we developed and tested mathematical models describing the kinetics of infection and immunity in anatomic compartments. Using these mathematical models, we also calculated the *R*_0_ of MCMV in the SG and predicted the probability of sustained viral replication and spread upon SG infection following different viral inoculation doses. Together, these results add to our understanding of the determinants of CMV infection and dissemination.

## Results

### Viral loads expand faster and decay slower in the SG than in other organs

The spread of MCMV using daily live luminescence bioimaging of mice following infection with a low dose of 10^3^ plaque-forming units (PFU; see Methods for dose determination) of a luciferase-tagged K181 strain of MCMV (K181-luc) to the right submandibular SG are shown in Fig 1. Virus was first noted solely at the site of inoculation (right submandibular SG), and then spread progressively throughout the body. Using two gates, we measured the strength of the luminescent signal in the SG compared to the rest of the body over time (Fig 2). Luminescence within the SG of infected mice was detectable and significantly higher (p-value <0.0005) than the background signal in uninfected mice as soon as 1 day post-infection. In the body, luminescence was not significantly greater in infected versus uninfected mice until 2 days post-infection (p-value <0.05). The total luminescent signal in the SG was greater than that seen in the body from days 5-21 post-infection despite the area of its gate being only 22% of the body’s. In both the SG and body, the signal rose quickly, peaking 7 days post-infection. Within the SG, the signal fit an exponential growth rate of 0.42/day, while the rate in the body was 0.14/day. After the peak 7 days post-infection, luminescence in the body declined markedly faster than in the SG, with fit exponential decay rates of 0.12/day and 0.03/day, respectively.

**Fig 1:**
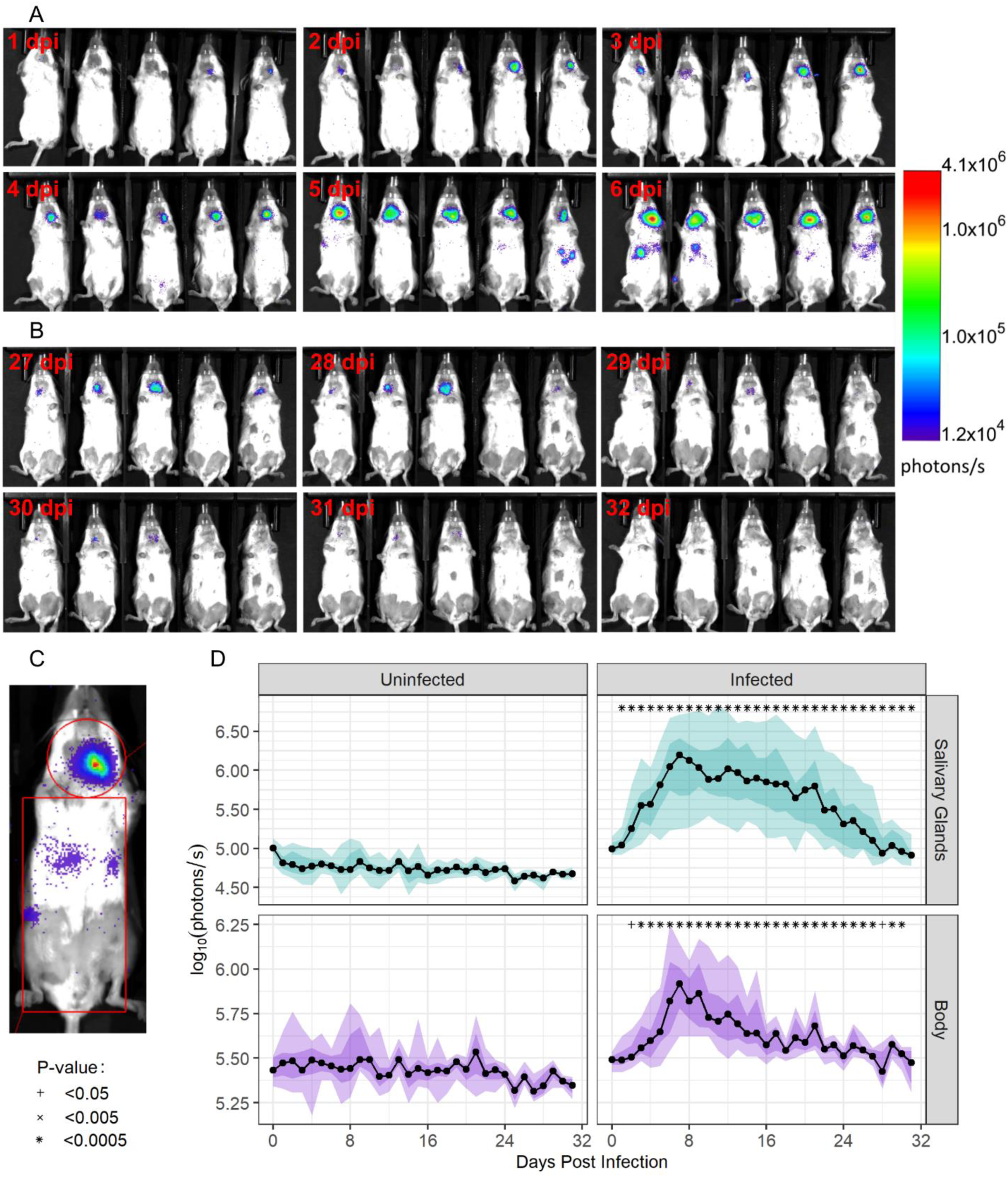
Spatiotemporal kinetics of viral MCMV dissemination from the SG. Bioimaging data from the first six days (**panel A**) and the last six days (**panel B**) post infection (dpi) are shown. Infection begins at the site of inoculation in the SG and disseminates throughout the body. Viral replication is greater in the SG and decays more slowly than in the rest of the body. By the end of observation (day 32), the signal within the SG has disappeared. The gates used to measure luminescent signal data in the SG separately from the other tissues (**panel C**). Longitudinal bioimaging data for these anatomical sites are shown for uninfected and infected mice (**panel D**). Symbols indicate the level of significant increase compared to background signal in uninfected mice on the same day.

**Fig 2:**
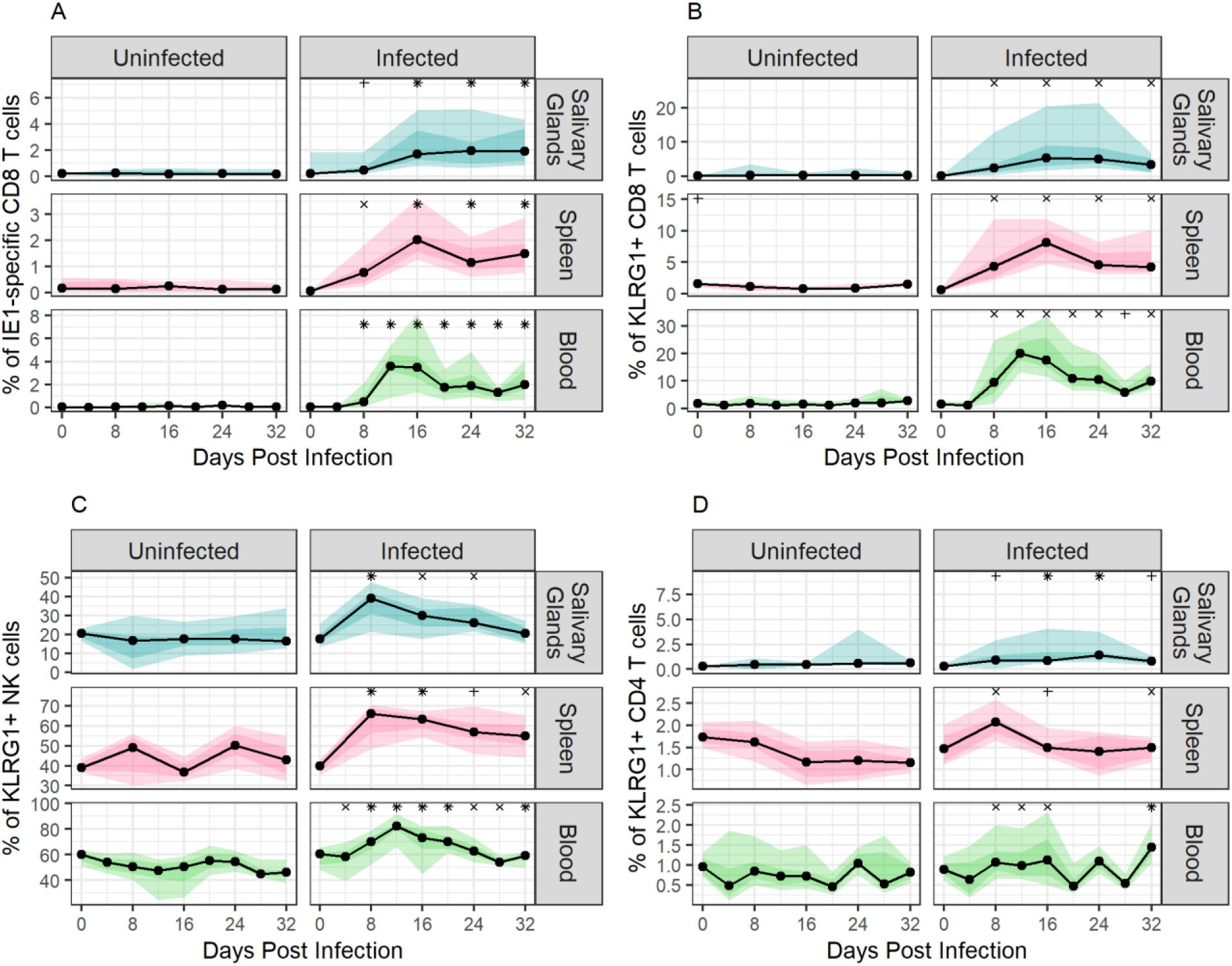
Expansion of immune cell populations during MCMV infection via the SG. Changes in immune cell populations within SG, spleen, and blood are shown: **panel A,** IE1-specific CD8 T cells; **panel B**, KLRG1+ CD8 T cells; **panel C**, KLRG1+ NK cells; **panel D**, KLRG1+ CD4 T cells. Immune cell population sizes are reported as the percentage of the parent population (CD8 T cells for panels A and B, NK cells for panel C, and CD4 T cells for panel D). Light ribbons show the 5-95% quantiles, dark ribbons show the 25-75% quantiles, black lines indicate median values, and dots indicate the time points at which data were collected. The symbols above the graphs indicate the level of significant increase compared to uninfected control values at the same time point, as defined in Fig 1.

### Subpopulations of CD8 T cells and NK cells, but not CD4 T cells, show significant changes throughout infection

Mononuclear cells isolated from whole blood, SG, and spleen were characterized by flow cytometry using markers to identify populations of B cells, NK cells, and CD8, CD4, and *γδ* T cells. To identify MCMV-specific CD8 cells, we included an MHC class I tetramer presenting the immunodominant IE1 epitope (15,19,34). We also stained for activation markers KLRG1, found on effector cells (15,35–37), and CD69, which has been associated with tissue-resident CD8 and CD4 T cells (29,37). Additional details are provided in the Methods section. The gating strategy used to identify cell populations of interest is shown in **Fig S. 1** of the Supporting Information.

Of the cell populations examined, IE1-specific CD8 T cells showed the most significant changes in size over time compared to those seen in uninfected mice (Fig 2A). These cells peaked in population size on days 12 and 16 post infection in the blood and spleen, respectively, while in the SG the population size plateaued on day 24 and was sustained until the end of the observation period. Large, significant changes were also observed in populations of KLRG1+ CD8 T cells, KLRG1+ NK cells, and KLRG1+ CD4 T cells in infected mice (Fig 2B-D, significance indicated). KLRG1+ CD8 T cells peaked between 12- and 16-days post-infection, depending on the site of collection, while KLRG1+ NK cells peaked between 8- and 12-days post-infection. KLRG1+ CD4 T cells peaked 8 days post-infection in spleen, 24 days post-infection in the SG, and 32 days post-infection in blood. These peaks in immune cell population sizes occurred a median of four days after the peaks in viral replication, as determined by the bioimaging signals. Flow cytometry data for other immune cell populations are shown in **Fig S. 2** of the Supporting Information. Smaller but statistically significant differences between uninfected and infected mice were noted for total populations of CD8 T cells, γδ T cells, and NK cells, consistent with previous findings that MCMV infection is primarily controlled by T cells and NK cells (15,29,38–40). There was no discernible change in total CD4 T cells or any CD69+ cell populations over the course of infection.

We next fit exponential growth models to the immune cell population dynamics in different tissues to compare the expansion rates before the peak was reached. During early infection, the frequency of IE1-specific CD8 T cells increased most rapidly in blood (rate of 0.338/day), followed by spleen (0.228/day), and SG (0.102/day). The frequency of KLRG1+ CD8 T cells increased at similar rates in all tissues (rate of 0.238/day in the SG, 0.191/day in blood, and 0.161/day in spleen). The rates of expansion of KLRG1+ NK cells were highest in the SG at 0.099/day, followed by spleen and blood with rates of 0.063/day and 0.018/day, respectively. Despite expanding fastest in SG, KLRG1+ NK cells represented a smaller proportion of the total NK cell population in the SG, being on average only 45.8% and 38.5% of those in the spleen and blood, respectively. The frequency of KLRG1+ CD4 T cells increased at a rate of 0.043/day in the spleen, 0.003/day in the blood, and 0.055/day in the SG.

### Mathematical models of MCMV infection

Few mathematical models of the within-host kinetics of HCMV infection have been published, and even fewer of MCMV infection (15,20,41). Based on the data we collected and information available in the literature, we created and fit two novel mathematical models to describe the dissemination of MCMV from its site of entry to the rest of the body, and to test which immune components are most important in controlling viral replication in each compartment.

#### Model 1: infection control by IE1-specific CD8 T cells

In our base model, we assumed that the observed large expansion of IE1-specific CD8 T cells is responsible for controlling infection in both the SG and the rest of the body. We supposed that MCMV in the SG and body (*V_b_* and *V_s_*, respectively) infects cells (*I_b_* and *I_s_*, respectively) at rates *η*_1_ and *η*_2_, respectively. As the virus infects a wide range of different cell types but does not impair organ function in this model (22), we assumed there is no target cell limitation. These infected cells produce MCMV at a per-capita rate of *p* and naturally die at a per-capita rate, *δ*. Infected cells stimulate the production of IE1-specific CD8 T cells (*T*) at a rate 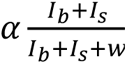, where *α* is the maximum proliferation rate and *w* is the number of infected cells needed for the proliferation rate to reach its half-maximum. In this model, we assumed that IE1-specific CD8 T cells target and kill both *I_s_* and *I_b_*, following the law of mass action, with rate constant *m*.

Upon ISG administration of MCMV we assumed that virus is present exclusively in the SG. Virus from the body and SG is assumed to disseminate to the other compartment at a per-capita rate *μ*. Equation set (1) shows all the ordinary differential equations (ODEs) for this model and a visual representation is provided in Fig 3A.

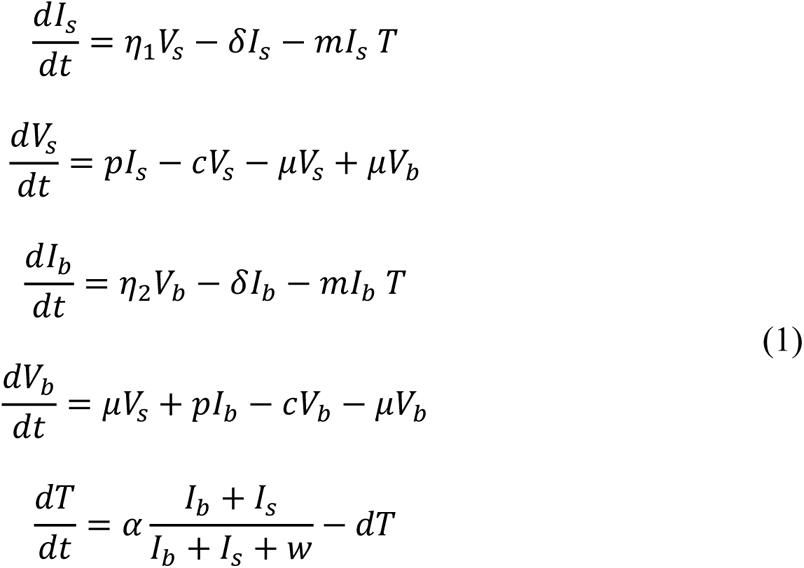

**Fig 3:**
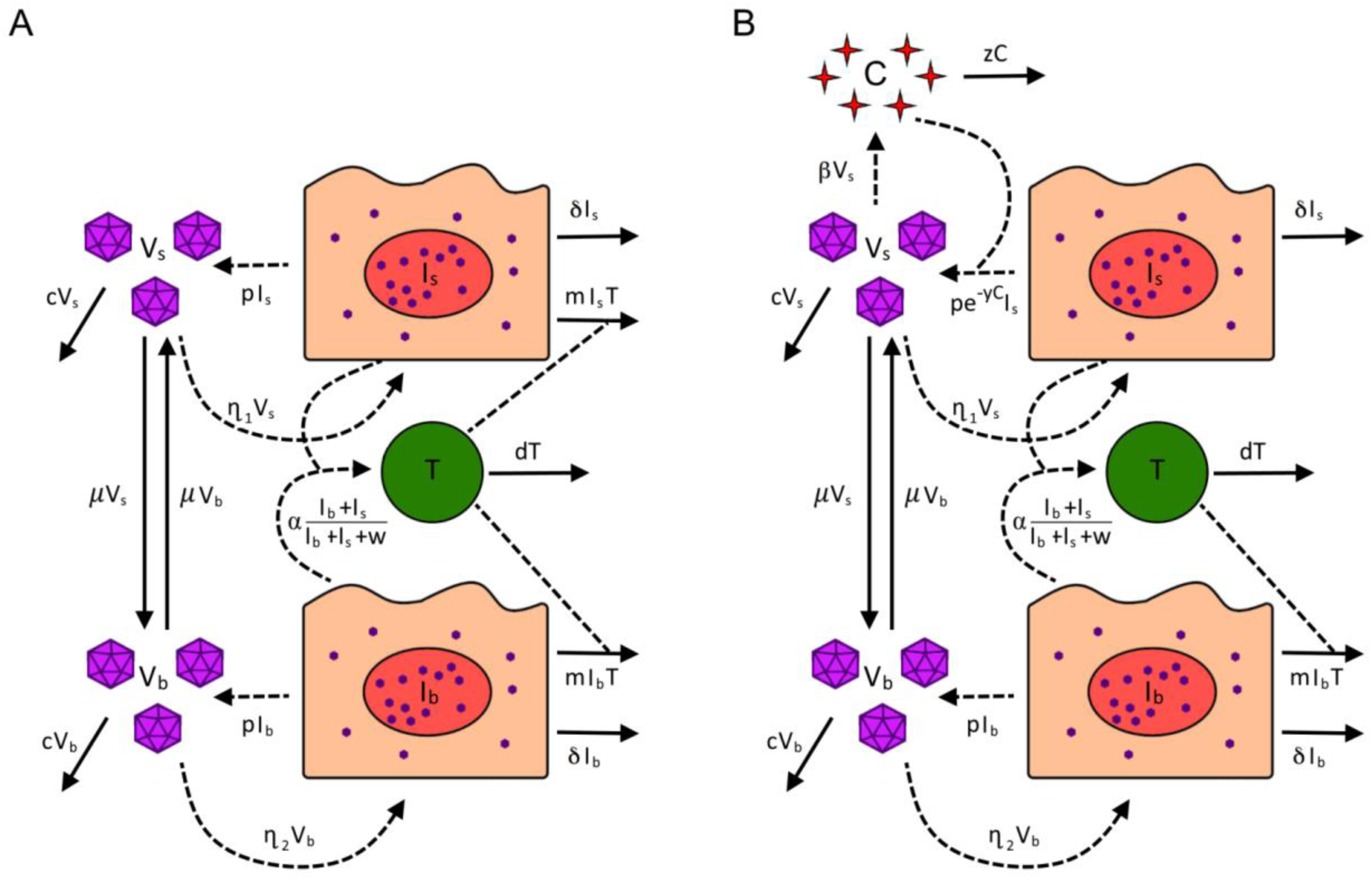
Visual representation of Models 1 and 2. In the body, infected cells (***I_b_***) are cleared by IE1-specific CD8 T cells (***T***). In Model 1 (panel A), infected cells in the SG (***I_s_***) are also cleared by IE1-specific CD8 T cells; however, in Model 2 (panel B), the production of virus in the SG (***V_s_***) is inhibited by IFN-***γ*** cytokines (***C***). Virus flows between the two compartments, allowing for the dissemination of infection.

#### Model 2: SG viral inhibition by cytokines

While we observed a large increase of IE1-specific CD8 T cells within the SG, MHC I expression has been found to be suppressed in MCMV-infected SG cells, thereby preventing their recognition and direct killing (30). However, significant expansion of activated CD4 T cells was also seen in the SG of infected mice (Fig 2D). As such, we developed a competing mathematical model consistent with elegant studies demonstrating that CD4 T cell-mediated cytokine release, principally IFN-*γ*, is critical for inhibiting MCMV replication in the SG (28,31–33). Our data and others suggest that this mechanism is far more important in the SG than in other parts of the body (30), where we found a less pronounced expansion of activated CD4 T cells compared to activated CD8 or NK cells over the course of infection.

To incorporate this immunological mechanism into the model, we supposed that cytokine production (*C*) occurs at a rate *βV_s_* in the SG. Due to suppression of MHC I expression on infected SG cells (30), we also assumed that these cells (*I_s_*) are no longer targeted by CD8 T cells (*T*) and, instead, cytokines inhibit viral reproduction in infected SG cells with an efficacy of *e^−yC^*. Cytokines in the SG decay at a rate, *z*. As the literature does not point to a direct role of CD4 T cells in controlling MCMV infection elsewhere in the body, the model assumes this effect is limited to the SG. Equation set (2) shows the full set of ODEs for Model 2, while a visual representation is shown in Fig 3B.

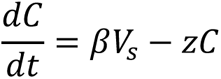

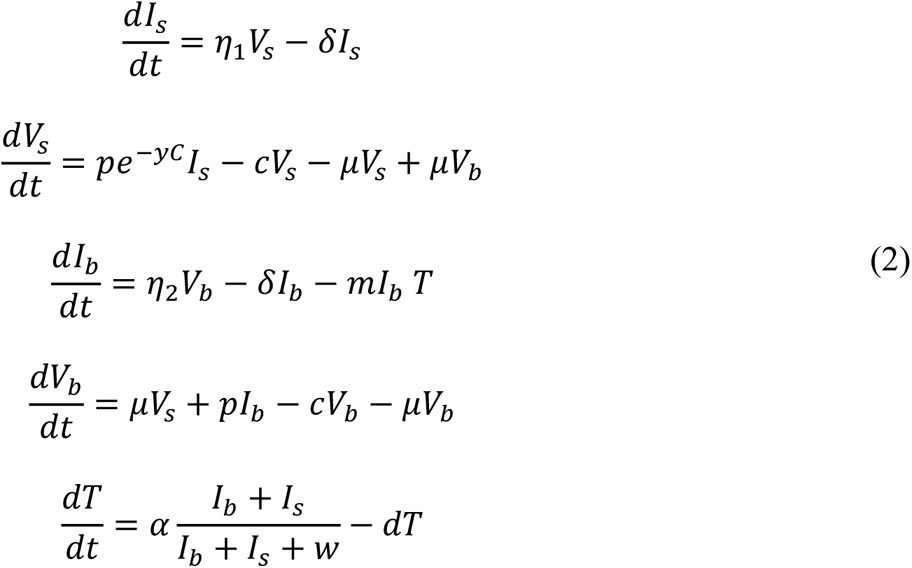

### CD8 T cell killing of infected cells does not explain the control of MCMV replication in the SG

We fit each mathematical model to pooled data from 10 ISG infected mice over 32 days post-infection, to test how well each model describes the data. Specifically, we fit *V_s_*to bioimaging signals in the SG, *V_b_* to bioimaging signals in the body, and *T* to the size of the IE1-specific CD8 T cell population in the blood (see the Methods section for details). We specifically used data from blood to fit *T* as were able to collect frequent longitudinal blood samples from mice, unlike from spleen or SG. During fitting, parameters with known values in the literature, or those that could not be distinguished during fitting, were left fixed, while others were allowed to vary. As such, parameters *m*, *α*, *μ*, *μ*, *η*_1_, *η*_2_, *y*, *β*, and *w* were fit while *z*, *p*, *δ*, and *c* were kept constant. Results of these fits are shown in Fig 4. The two model fits were compared using the Akaike information criterion (AIC), which evaluates the prediction error of each model. Consistent with experimental observations (20,30–32), Model 2 (CD4 T cell-derived IFN-*γ*) outperformed

**Fig 4:**
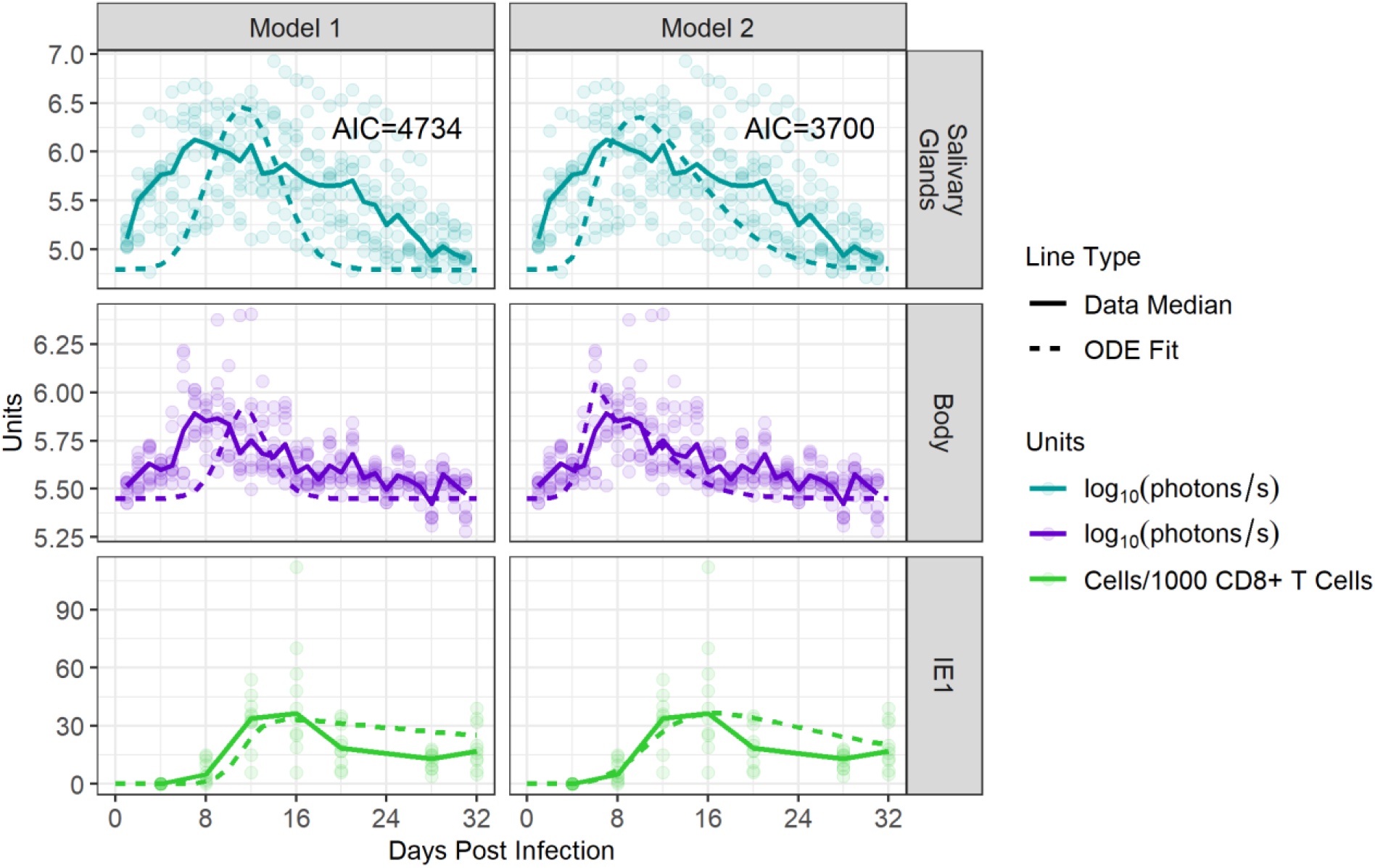
Control of viral replication in the SG is better explained by CD4 T cell-mediated cytokine production than direct killing by CD8 T cells. We compared how well each mathematical model was able to reproduce the observed murine data. Simultaneous fits for each model across 10 mice are shown. Dots represent luminescent signals captured in the SG and body during bioimaging and the number of IE1-specific CD8 T cells/1000 CD8 T cells within the blood. Solid lines indicate median values. Dotted lines show the optimal ODE fit, as determined by our fitting algorithm. AIC values for each model are shown.

Model 1 (direct killing by IE1-specific CD8 T cells) with a ΔAIC of 1034. With such a large ΔAIC, these results indicate that Model 2 better explains the data and that control of salivary gland infection is attributable more to cytokines, rather than to IE1-specific T cells as in Model 1. In particular, Model 2 better captured the fast rise in viral load (VL) observed in experiments. Thus, all further data analyses were performed using Model 2.

We next fit Model 2 to data from each infected mouse to arrive at one set of best-fitting parameter values for each animal. Examples of individual fits are shown in Fig 5A, and the general trend seen over time for all model compartments is shown in Fig 5B. Remaining fits for other ISG-infected mice are shown in **Fig S. 3** of the Supporting Information. The median value and 5-95% quantiles for each fit parameter when pooling all fits are shown in

**Table 1:** Parameters used in the mathematical model. Numbers marked with a (*) indicate parameters that were estimated by fitting Model 2 to data. (+) indicates the number was estimated based on values in the literature to determine the best value to match the kinetics of infection and kept constant during fitting.

| Parameter | Description | Units | Literature Values | Estimate |
| --- | --- | --- | --- | --- |
| $m$ | Rate at which $T$ kills $I_b$ via mass action | $day^{-1}$ | 0.01<br>(42) | $6.33 \times 10^{-1*}$<br>($1.01 \times 10^{-4}$, 1) |
| $\alpha$ | Maximum rate at which $I_b$ and $I_s$ stimulate production of $T$ | $day^{-1}$ | — | $1.93 \times 10^{2*}$<br>($9.49, 2.33 \times 10^4$) |
| $d$ | Death rate of $T$ | $day^{-1}$ | 0.05-0.322<br>(15,42) | $8.38 \times 10^{-2*}$<br>($1.02 \times 10^{-2}$, $9.91 \times 10^{-1}$) |
| $\mu$ | Rate of viral exchange between SG and body | $day^{-1}$ | — | $5.33 \times 10^{-1*}$<br>($6.40 \times 10^{-4}$, 8.32) |
| $\eta_1$ | Rate at which $V_s$ causes new cellular infection | $day^{-1}$ | 0.6<br>(42) | $2.61 \times 10^{-1*}$<br>($1.97 \times 10^{-1}$, $4.38 \times 10^{-1}$) |
| $\eta_2$ | Rate at which $V_b$ causes new cellular infection | $day^{-1}$ | 0.6<br>(42) | $5.74 \times 10^{-2*}$<br>($1.00 \times 10^{-3}$, $3.24 \times 10^{-1}$) |
| $y$ | Exponential rate at which $C$ inhibits the production of $V_s$ | $cytokine^{-1}$ | — | $5.32 \times 10^{-5*}$<br>($6.50 \times 10^{-8}$, $6.10 \times 10^{-4}$) |
| $\beta$ | Rate at which $V_s$ stimulates the production of $C$ | $day^{-1}$ | — | $2.05 \times 10^{-3*}$<br>($2.05 \times 10^{-3}$, 4.02) |
| $w$ | Number of infected cells needed for T cell production to reach its half-max rate | $cells$ | — | $1.21 \times 10^{7*}$<br>($7.27 \times 10^5$, $9.15 \times 10^8$) |
| $z$ | Decay rate of $C$ | $day^{-1}$ | 3.6(20) | 0.01<sup>+</sup> |
| $p$ | Production rate of viruses by infected cells | $day^{-1}$ | 9.84-1600(20,33) | 100<sup>+</sup> |
| $\delta$ | Natural death rate of infected cells | $day^{-1}$ | 0.77-1.2(33,42) | 1<sup>+</sup> |
| $c$ | Decay rate of viruses | $day^{-1}$ | 2-10.8<br>(20,33) | 8.8<sup>+</sup> |
| | Mean bioimaging background signal from bioimaging | $photons/s/cm^2$<br>$/steradian$ | | $1.57 \times 10^3$ |
| | Bioimaging SG gating area | $cm^2$ | | 3.13 |
| | Bioimaging body gating area | $cm^2$ | | 14.2 |

**Fig 5:**
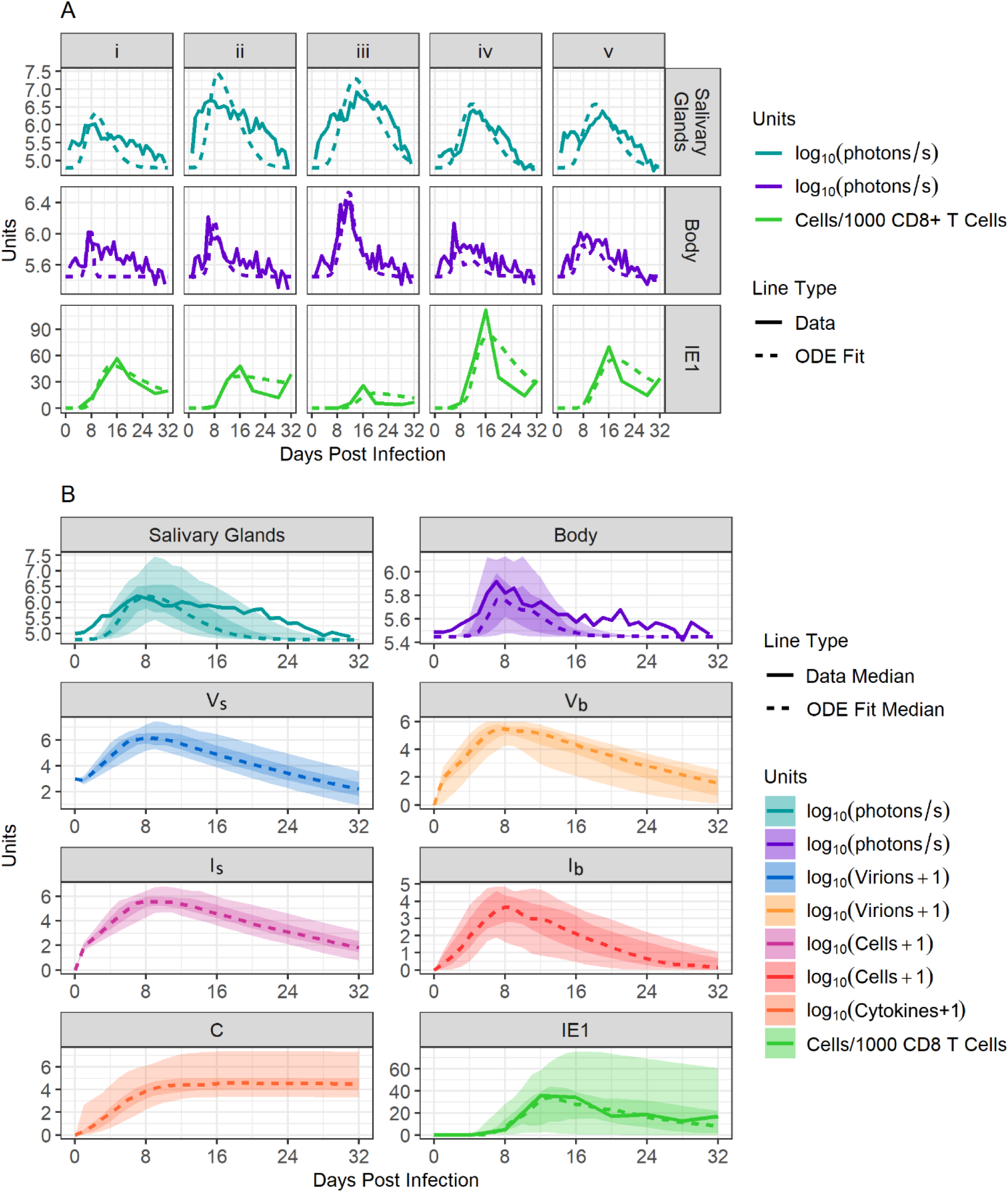
Mathematical modelling of primary MCMV infection. **Panel A**: Model 2 fit, with data from 5 mice separately. **Panel B**: Summary of fits for all mice and for all compartments of the model. Dotted lines show the median value of best fitting simulations, while solid lines show the median value of collected data (when a comparison was available). Dark ribbons show the 25-75% quantiles and light ribbons show the 5-95% quantiles.

Having generated estimates of all parameter values in our model, we next compared how parameter values governing the infection dynamics within the SG and the rest of the body differ and estimated how quickly MCMV is exchanged between these compartments. Our model predicts that the rate of infection within the SG *η*_1_, is significantly faster than the rate of infection within the body, *η*_2_, (p-value<0.05) coinciding with the high luminescence signals observed in the SG. We also noted that the exchange of virus between the body and SG is quite fast, occurring at a median rate of 0.553/day, which corresponds to a half-life of residency of approximately 30 hours.

We also found that while IE1-specific CD8 T cells, which control infection within the body, decay at a median rate of 0.08/day, cytokines controlling infection within the SG were fit to a slower decay rate of 0.01/day. This slower decay rate indicates that cytokine levels are maintained for a long period (Fig 2C), causing sustained suppression of viral replication in these glands. We found that faster decay rates of cytokines led to oscillating VL that were not observed biologically (results not shown).

### Mathematical modelling predicts a high within-host basic reproductive number for MCMV

Using the estimated parameter values, we calculated the within-host basic reproductive number (*R*_0_) for MCMV in the SG. Here, *R*_0_is defined as the number of infected cells propagated by a single infected cell in the absence of any immunity. For our mathematical model, *R*_0_ is defined as the dominant eigenvalue of the model’s next generation matrix (43), and equals

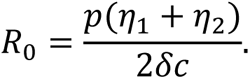

Calculating *R*_0_using our fit parameter values gave a median *R*_0_ value of 2.2 (5-95% quantiles of 1.5-3.5). As a point of comparison, the within-host infection *R*_0_ value was estimated to be 1.6 for HCMV using clinical data obtained during infant primary infections (24).

### Low-dose primary SG infections in mice are predicted to persist and spread

To conclude our mathematical analysis of MCMV dynamics in the SG, we used our model to predict the relationship between the ISG inoculum and viral spread. By simulating the stochastic analogue of the system of ODEs described in Model 2, and using parameter values obtained through fitting (), we varied the initial dose assumed to be injected into the SG. Though this analysis, we identified which inoculation doses are predicted to result in persistent SG replication and systemic dissemination, and which inoculations may cause brief self-limited SG infection. Results are shown in Fig 6A.

**Fig 6:**
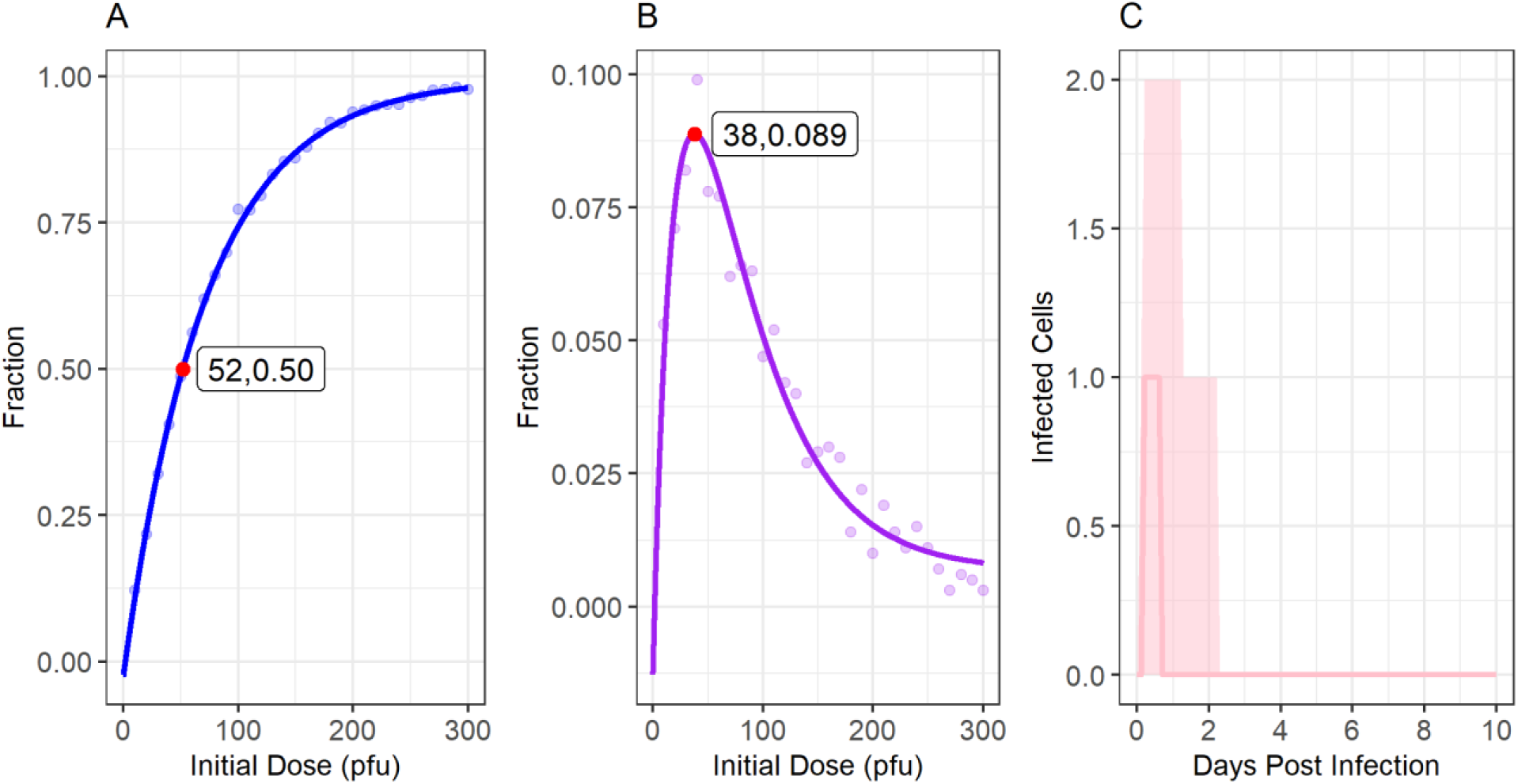
Modelled spread of SG infections in mice. **Panel A**: We modelled the fraction of SG infections that disseminate beyond the SG in mice as a function of the initial ISG dose. The red dot shows that our model predicts the ***ID*_50_**, the ISG dose at which 50% of infections spread beyond the SG, to be 52 PFU. **Panel B**: The fraction of inoculations that cause transient local infection in the SG as a function of the initial dose. Here, a transient infection is one that infects SG cells but dies out before spreading to the body. As indicated by the red dot, our model predicts transient infection is most likely with an initial dose of 38 PFU, occurring after 8.9% of inoculations. **Panel C**: Our model’s predictions on the number of infected cells among infections that are limited to the SG over time when inoculating mice with an ISG dose of 38 PFU. Among infections that do not disseminate, very few cells become infected (median maximum of 1 cell, 5-95% quantiles of 1-3 cells), and replication dies out very quickly, taking a median of 0.7 days (5-95% quantiles of 0.3-2.1 days) to be cleared. Lines in panels A and B show the line of best fit. The line in panel C indicates the median behaviour, and light ribbons show the 5-95% quantiles over time.

Our model predicts that with a dose of 52 PFU of K181-luc administered ISG, 50% of mice will have a sustained infection that disseminates throughout the body (*ID*_50_; Fig 6A). These results are supported by our findings that no mice were infected at a dose of 10 PFU via the SG, but approximately two-thirds of mice get infected at a dose of 100 PFU (results not shown). At doses of 300 PFU, and 500 PFU, our model predicts that 98% and 100% of mice, respectively, would have a systemic infection.

Our model also predicts that transient SG infection, with limited viral replication within the SG that dies out before spreading to the rest of the body (Fig 6B-C) is possible with low-PFU inoculations. However, transient infections are still predicted to be rare and, when occurring, a median of only 1 cell (5-95% quantile of 1-3 cells) within the SG is predicted to be infected at any time. These infections are also predicted to die out very quickly, only lasting a median of 2 days (5-95% quantile of 2-4 days). This phenomenon is likely due to the predicted high rate of viral exchange between the SG and the rest of the body (*μ*) and a relatively high *R*_0_value, suggesting that once cells are infected in the SG, replication almost always persists, and typically also spreads rapidly to the rest of the body.

### Fitting our mathematical model to other MCMV infection data

To validate our model, we next examined whether infections via the IP route with different inocula of MCMV were consistent with Model 2. Mice were infected with either a low (10^2^PFU) or a high (10^6^ PFU) dose of K181-luc, imaged daily for luminescence, and blood samples were taken every seven days to measure changes in immune cell populations. Model 2 fit these new data well, reproducing the rise and fall in VL and immune cell population sizes. Data and fits from mice infected with 10^2^PFU IP and 10^6^PFU IP are shown in **Fig S. 4** of the Supporting Information.

Finally, we looked at how the parameter values predicted when fitting Model 2 to data from ISG inoculation versus IP inoculation compared. Distributions of fit parameters for each data set are shown in Fig 7. In general, estimated parameter values were similar with different ROA. Values for *η*_1_, *m*, *d*, and *μ* showed small but significant differences across data sets (Fig 7). The largest most significant differences were seen for parameter *η*_2_, which was estimated to be significantly larger when fitting the model to data from IP infected mice than when fitting it to data from ISG infected mice.

**Fig 7:**
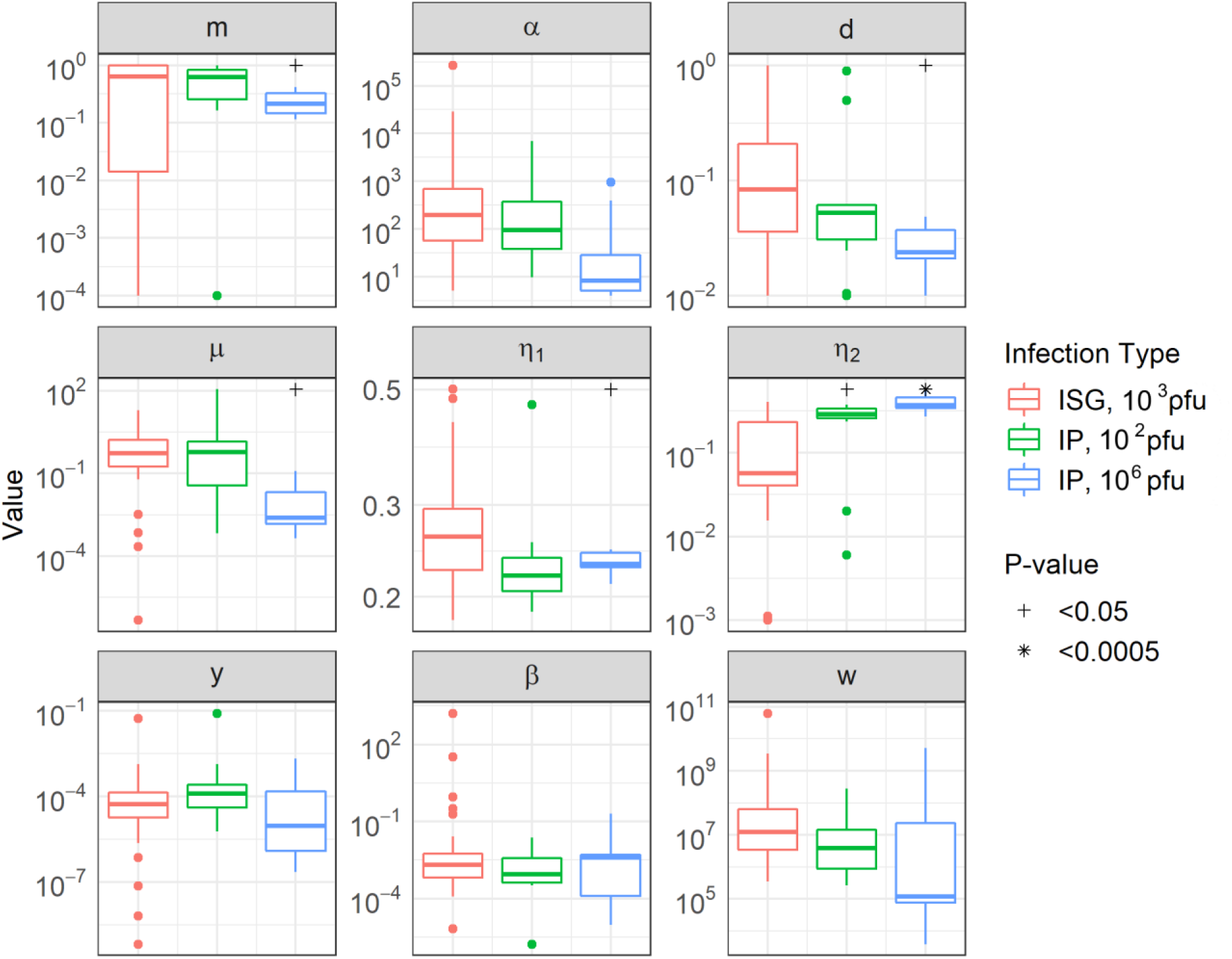
Parameter distributions for model fit parameters when fitting individual mouse data. Parameter distributions across the data sets were stratified to fit Model 2. Significant differences were seen between the “fit” of parameter values using ISG-infected mice and their fit using IP-infected mice.

## Discussion

A deeper understanding of the kinetics and immune correlates of CMV SG replication has the potential to inform the design of vaccines to prevent infection and transmission. Through collecting comprehensive time-series data following a low dose ISG infection of MCMV in mice, we identified organ-specific fluctuations in key immune cell populations and their temporal relation to viral replication dynamics. Using these experimental data, we designed and fitted novel mathematical models describing the spatial spread of MCMV and the immune responses within different compartments of the body to glean insight into the determinants of systemic infection and immune control.

IE1-specific CD8 T cells expanded at the highest rate following infection. However, lasting and significant elevations in populations of KLRG1+ CD8 T cells, KLRG1+ NK cells, and KLRG1+ CD4 T cells were also observed, eventually contracting with decreasing viral replication. We anticipated differences in immune cell dynamics according to anatomic compartment given the relatively greater and longer viral replication in SG. Indeed, virus luminescence rose three times faster during the early stages of infection and declined four times slower following signal peak in SG than the rest of the body. While weaker IE1-specific CD8 T cell and KLRG1+ NK cell responses were observed in SG than at other sites, all four immune cell populations generally displayed similar kinetics in all compartments. This suggests that despite the presence of similar immune cell populations at different anatomic sites, their ability to recognize and eliminate infected cells differs. In support of other studies (20,30–32), our mathematical analysis suggested that killing of infected cells by virus-specific CD8 T cell is sufficient to explain viral kinetics only outside the SG. In contrast, the model requires cytokine production by CD4 T cells in the SG to accurately reproduce the experimental data.

Our mouse model used small amounts of virus delivered via ISG in an attempt to mimic human infection, which allowed us to characterize the rate of persistence and spread within and beyond the SG. Oral HCMV infection may at times die out before causing a full systemic infection, based on prospective cohort data, in which brief, low-level episodes of viral shedding in saliva can be observed in individuals in the absence of seroconversion (10,33,44,45). Self-limited local infections appear to be due to a low within-host *R*_0_for HCMV, estimated at 1.6 in the infant oral cavity and thus quite poor cell-to-cell spread of infection in the oral mucosal epithelium (33). In contrast, our mathematical model estimates an *R*_0_ of 2.2 for MCMV in the SG of our experimental animals. Further, while previous research has suggested that ISG ROA of MCMV leads to reduced systemic pathology as compared to other ROAs (13), our model suggested viral spread from the SG to the rest of the body is still quick and efficient, such that self-limited SG infections are rare and last only 1-2 days.

The observation that MCMV disseminates more efficiently than HCMV may simply represent intrinsic differences in these viruses, given that MCMV replication lasts days-weeks after primary infection compared to weeks-months for HCMV (24) Importantly, the efficiency of viral spread measured using the MCMV strain K181, which is highly laboratory adapted, may not reflect wild-type strains. Further, we cannot rule out the possibility that direct injection into mouse SG tissue in the mouse differs from natural oral HCMV acquisition. For example, trauma resulting from ISG inoculation could have could favour faster spread to other anatomic sites. In addition, other oral epithelial cell types may be infected prior to SG in humans. HCMV infection is often acquired early in life, through frequent, repeated exposures (46–48), as opposed to a single inoculation into the SG. Breast milk, a common source of infection in infants, also contains a host of antibodies and other immune factors that may influence the likelihood and course of infection (49,50). Further, while the SG is indisputably a site of early viral infection in both humans and mice (14,16,18), elegant studies indicate that natural infection in the mouse is likely acquired through the nose (17,23,51). Thus, future models should be informed by experimental infections employing intranasal inoculation or breast milk transmission.

Our results also bear significant relevance for the design of vaccines aimed at preventing infection or minimizing shedding (10,52), and thereby curbing transmission to pregnant women, an approach that appears highly effective in preventing cCMV (53–55). By revealing the unique persistence of viral replication within the salivary glands despite the presence of similar infection-induced immune cells to those observed in the rest of the body, our findings underscore a critical point: the requirements for a vaccine to confer protection or minimize shedding in the salivary glands likely differ significantly from those needed at other bodily sites. With the probable importance of the salivary glands in oral transmission, both as a site of initial exposure and as a contributor to the amount of virus shed into saliva, this aspect may become a crucial component in the design of a successful vaccine. Consequently, vaccine strategies emphasizing the stimulation of IFN-γ and TNF-α, which appear necessary for salivary gland CMV control, rather than simply a robust CD8 T cell response, may emerge as essential requirements for preventing or mitigating the duration and severity of infection.

## Materials and Methods

### Virus and inoculation of mice

Female BALB/c mice obtained from Charles River were infected with a variant of the K181 strain of MCMV with the *m78* gene tagged with luciferase (generously gifted by Helen Farrell, University of Queensland). A full description of this construct has been described elsewhere (18). Virus stocks were grown in M2-10B4 cells (ATCC # CRL-1972) with RPMI 1640 Medium special formulation (Thermo Fisher cat # A1049101) supplemented with 10% fetal bovine serum (Thermo Fisher cat # 12483020) and 1% penicillin-streptomycin (Thermo Fisher cat # 15140148). Mice were infected via ISG or IP administration. For ISG administration, a 5 *μ*l solution containing 1000 PFU of K181-luc and PBS was prepared and injected with a syringe directly into the right submandibular SG while the mouse was under isoflurane anasthesia. Preliminary tests performed indicated this to be the lowest dose necessary to ensure infection of all mice following ISG inoculation (data not shown). For IP inoculation, a 100 *μ*l solution containing either 10^2^ PFU or 10^6^PFU of K181-luc was diluted in PBS and injected with a syringe directly into the peritoneum of mice while they were awake and scruffed. All mice were between the ages of 6 and 10 weeks when inoculated. A total 39 mice were infected ISG with 1000 PFU, 11 mice were infected IP with 100 PFU, and 11 mice were infected IP with 10^6^ PFU. For every infected mouse, a control mouse was administered PBS, either ISG or IP, and monitored at the same time and treated in the same way as infected mice.

### Bioimaging

Mice received an IP injection of 100 *μ*l of a 2% D-luciferin solution (Goldbio cat #115144-35-9), were anaesthetized with isoflurane gas, and transferred to a Spectral Instruments Ami HTX bioimager for monitoring of light emission with a CCD camera. Bioimaging data was analyzed using the Aura Image Analysis software.

### Tissue and blood sample collection and flow cytometry

Blood was collected from mice via the saphenous vein every four days for mice infected via ISG administration, and every seven days for mice infected via IP administration. Spleens and SG were harvested every eight days from subsets of ISG infected mice. Spleens were homogenized and strained through a 70 *μm* mesh to yield a single-cell suspension. SG were processed using the MACS Miltenyi multi-tissue dissociation kit (order no. <u>130-110-201</u>) to create a single-cell suspension. Blood and spleen cell suspensions were further incubated with an RBC lysis buffer (eBioscience, cat # 00-4300-54). Single-cell suspensions were then stained with eFluor 780-conjugated viability dye (Invitrogen eBioscience cat # 65-0865-14), and fluorescently tagged with monoclonal antibodies against CD3 (PerCP-eFluor 710, eBioscience cat # 46-0032-82), CD19 (BV-510, BioLegend, cat # 115545), CD4 (BV-785, BioLegend cat # 100453), CD8a (BUV-737, BD Bioscience cat # 564297), gd (BUV-563, BD Bioscience cat # 748993), CD69 (PE-CF594, BD Bioscience cat # 562455), KLRG1 (APC, BioLegend cat # 138411), CD335 (BV-711, BD Bioscience cat # 740822), CD49b (PE-Cyanine7, eBioscience cat # 12-5971-82), and MHC class I tetramer containing the FITC-labelled H-2L^d^ 168-YPHFMPTNL-176 peptide produced by the *ie1* MCMV gene (obtained from the NIH Tetramer facility core). Cells were analyzed for the presence of fluorophores using the BD FACSymphony™ flow cytometer. Flow cytometry data was analyzed and gated using FlowJo software.

### Statistical Analysis

Statistical significance of differences between data from infected and uninfected mice at specific time points was determined using the Mann-Whitney test. *P*-values less than 0.05 were considered statistically significant. Rates of exponential growth and decay of immune cell populations and luminescent signals were analysed by fitting a linear model to the number of days post-infection and the log-transformed data. For exponential growth, only data points collected before the median peak value were included. For exponential decay, only data points collected after the median peak value were included.

### Model simulation and parameter estimation

Mathematical models were simulated using the R package, “pomp” (56). Parameters of the model were fit by matching the trajectories of the deterministic model to our data. Here, we chose distributions to determine the probability of model predictions given the observed data and used these to create a likelihood function. We then created an objective function meant to evaluate the likelihood function and used the Nelder-Mead method to search parameter space to find parameters that maximized this likelihood. Throughout fitting, we kept parameters *z*, *p*, *δ*, and *c* fixed while allowing all other parameters defined in the set of ODEs to vary.

### Defining the likelihood function

Let *V_b_*(*t*) be the model-predicted number of virions present in the body at time *t*, *a* be the measured number of photons/s released per virion, *B_b_* be the average background signal in the body as measured in uninfected mice, and *M_b_*(*t*) be the bioimaging signal measured in the body at time *t* in units of photons/s. We then assume *aV_b_*(*t*) + *B_b_* follows a lognormal distribution with mean *M_b_*(*t*) and standard deviation *ρ*_1_.

Similarly, letting *V_s_*(*t*) be the number of virions present in the SG at time *t*, *B_s_* be the average background signal in the salivary gland, and *M_s_*(*t*) be the bioimaging signal measured in the SG at time *t* in units of photons/s, we assume *aV_s_*(*t*) + *B_b_* follows a lognormal distribution with mean *M_s_*(*t*) and standard deviation *ρ*_1_.

For comparing model predicted numbers of IE1-specific CD8 T cells to data, we let *T*(t) be the model-predicted number of T cells in the blood at time *t*, *f* be the average number of CD8 T cells in the blood, *ρ*_2_ be a cell’s probability of being observed through flow cytometry, and *F_IE1_*(*t*) be the measured fraction of CD8 T cells that are IE1-specific in the blood at time *t*. Thus, we assume *T*(*t*) follows a Poisson distribution with rate *ρ*_2_*fF_IE1_*(*t*).

With these assumptions, we define the likelihood function as

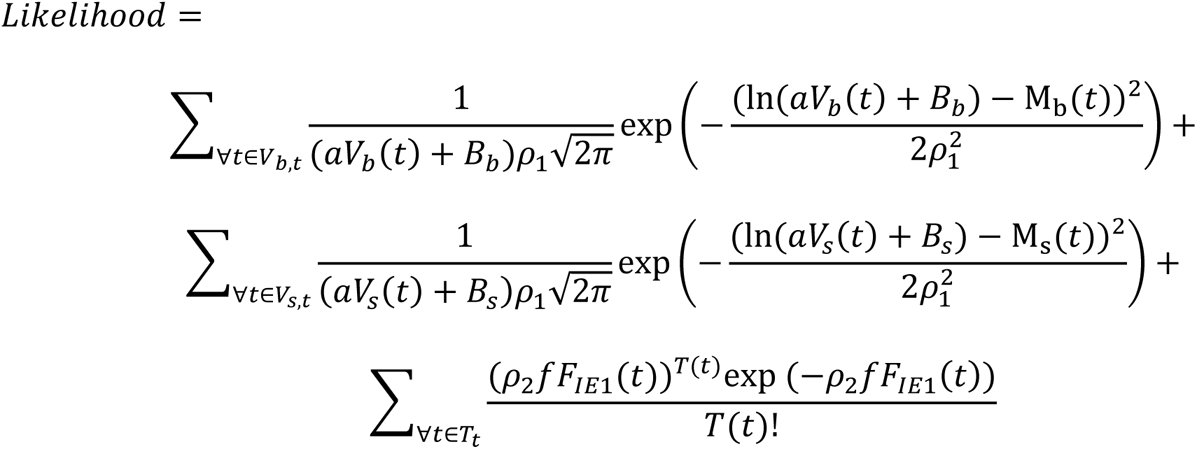

where *V_b,t_* is the set of times where *M_b_* was measured, *V_s,t_* is the set of times where *M_s_* was measured and *T_t_* is the set of times *F_IE1_* was measured.

### Stochastic Simulations

Stochastic simulations of the model were performed by converting the deterministic skeleton of the mathematical model into a series of individual reactions. The model progresses through time following the tau-leaping algorithm where small time steps of 0.001 days were made (57). At each time step, the number and type of reactions occurring were randomly chosen from a Poisson or Multinomial distribution, depending on the independence of the reaction, with the probability dependent on the reaction rate.

## Supporting Information

**Fig S. 1:**
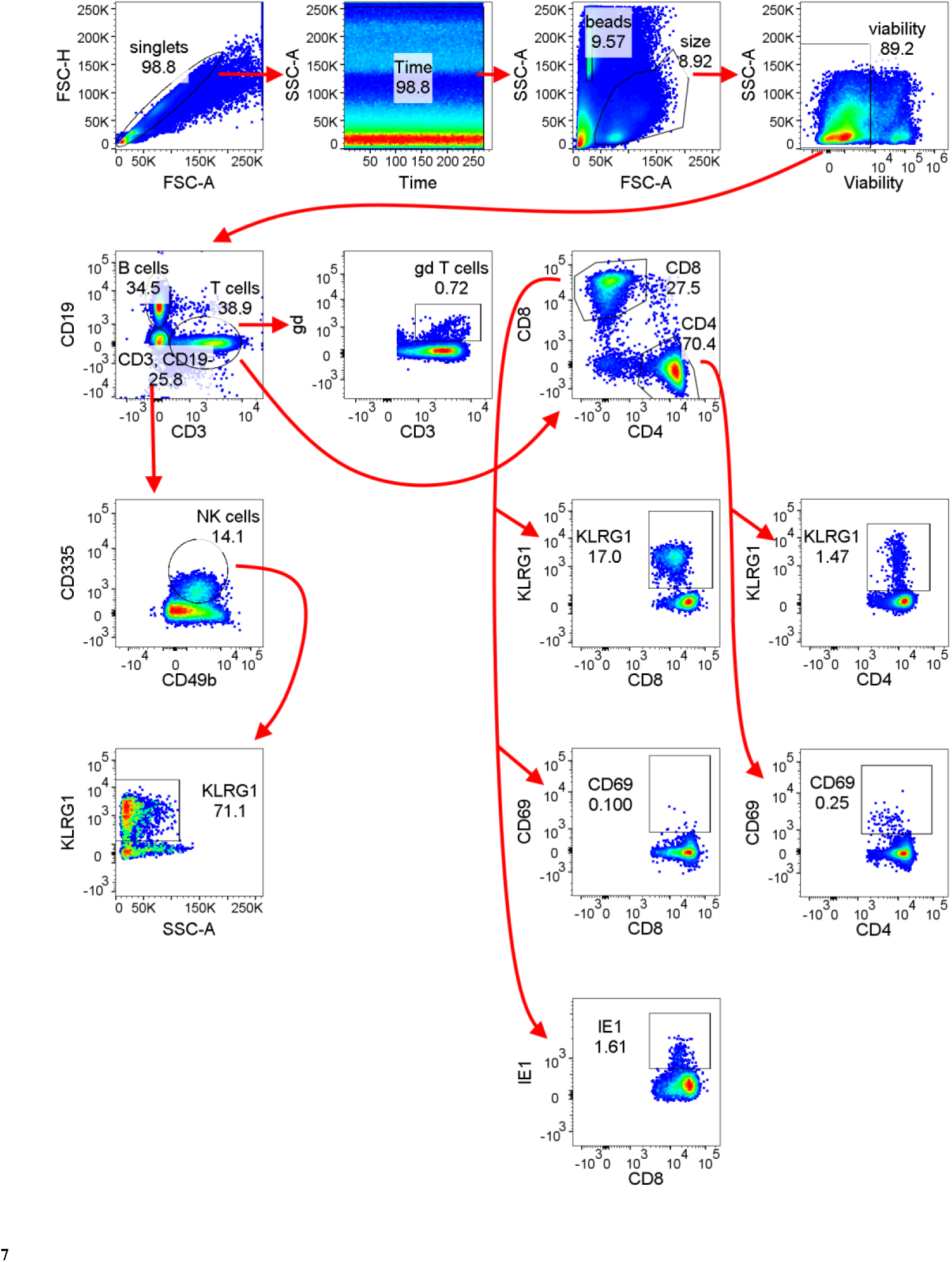
Gating strategy used to identify immune cell populations of interest. Cells were first gated against FSC-H and FSC-A to remove doublets, then against time and SSC-A to ensure no acquisition issues. We further gated against FSC-A and SSC-A to identify cells of the appropriate size, and against SSC-A and the viability dye used to identify live cells. Live cells were then gated using remaining markers to identify the cell populations of interest.

**Fig S. 2:**
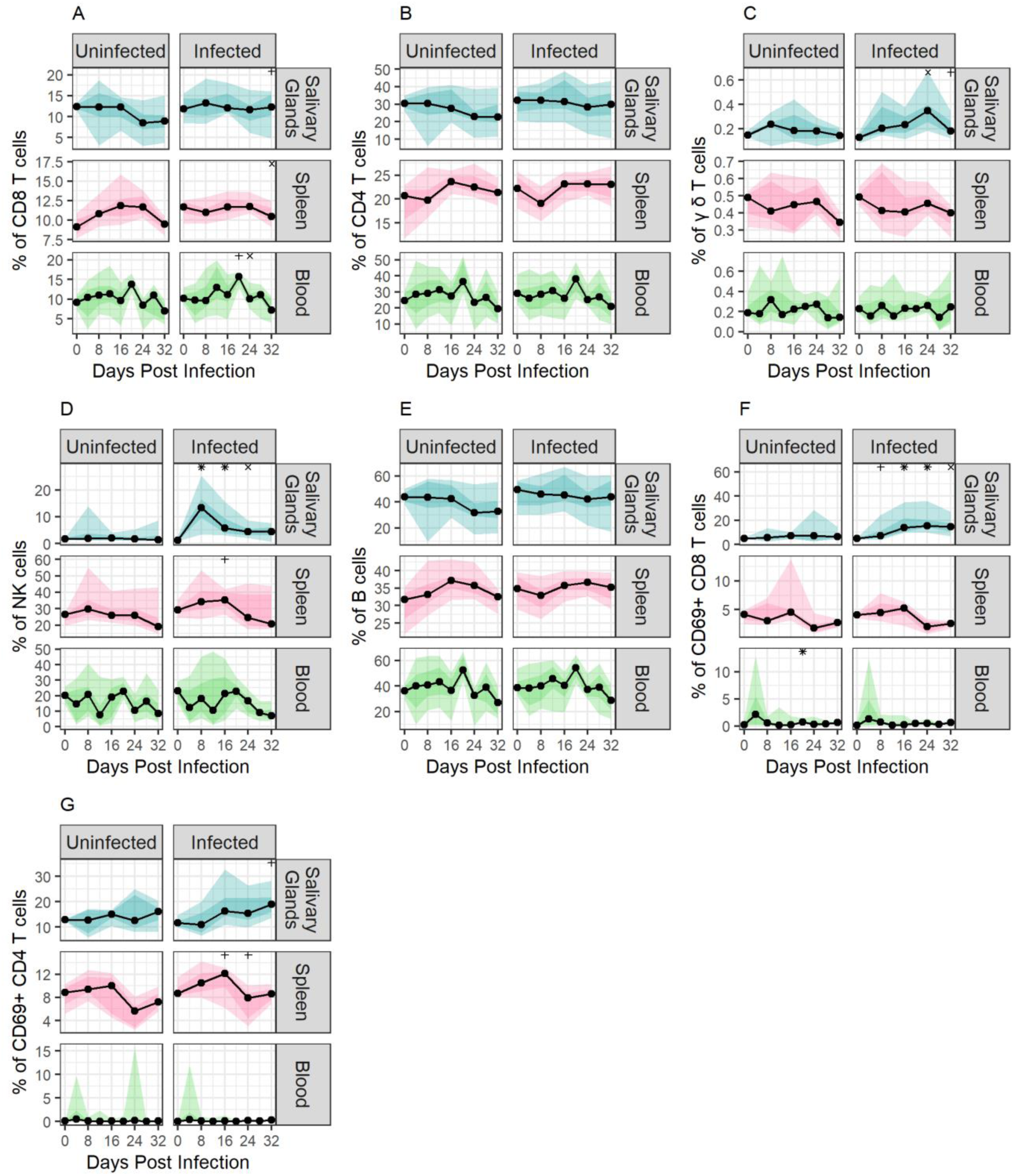
Immune cell populations of secondary interest and their change over the course of observation in uninfected and MCMV-infected mice. Symbols +, ×, and * above data indicate days where an immune cell proportion was significantly different between uninfected and infected mice. Symbol “+” represents where the p-value was less than 0.05, symbol “×” represents where the p-value was less than 0.005, and symbol “*” represents where the p-value was less than 0.0005. The symbol position is always above the group that had a higher median value than its comparator. Plots A-C are reported as the percentage of viable cells while D-G are reported as percentage of parent population.

**Fig S. 3:**
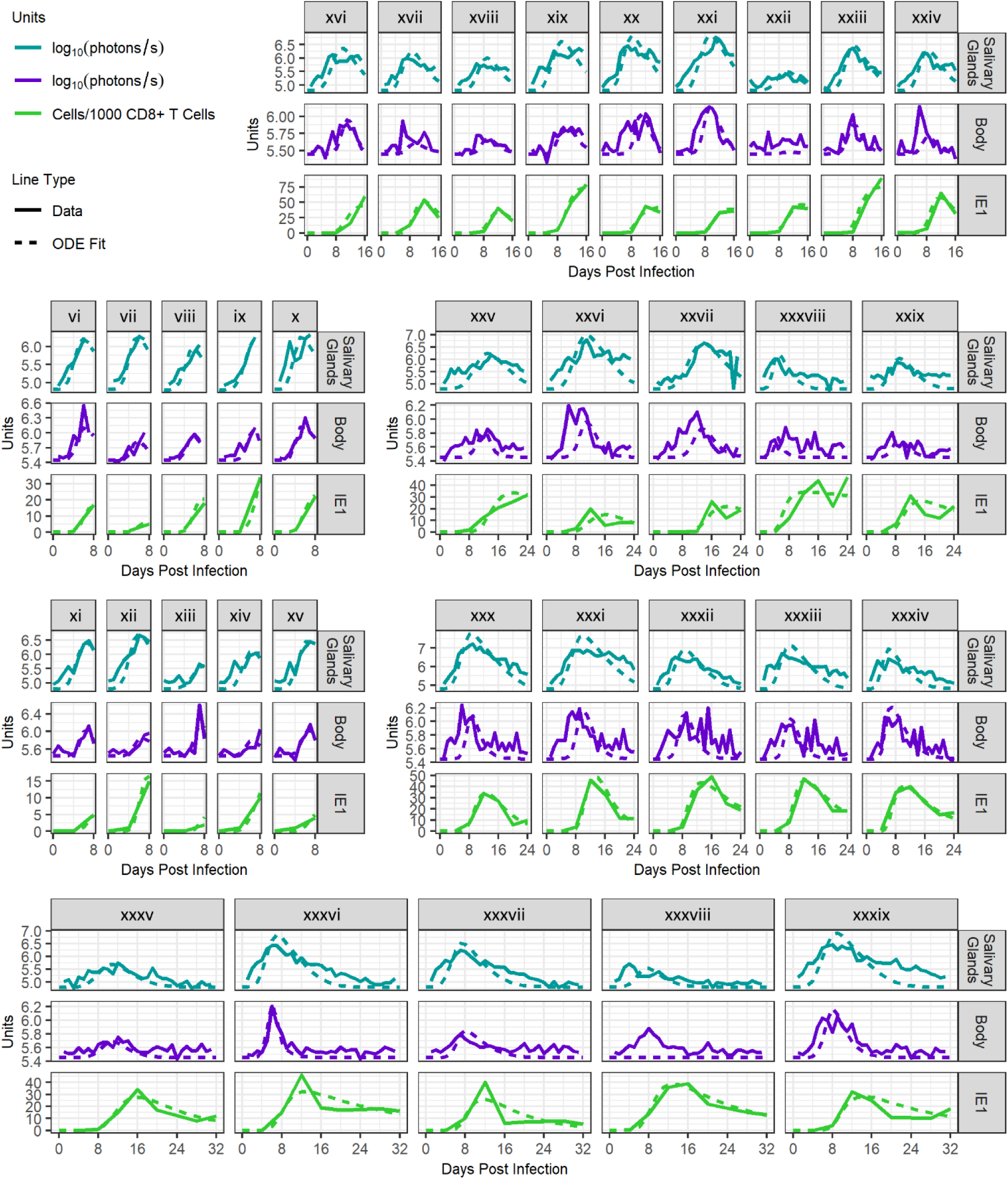
Additional fits to mice infected ISG with 1000 PFU K181-luc.

**Fig S. 4:**
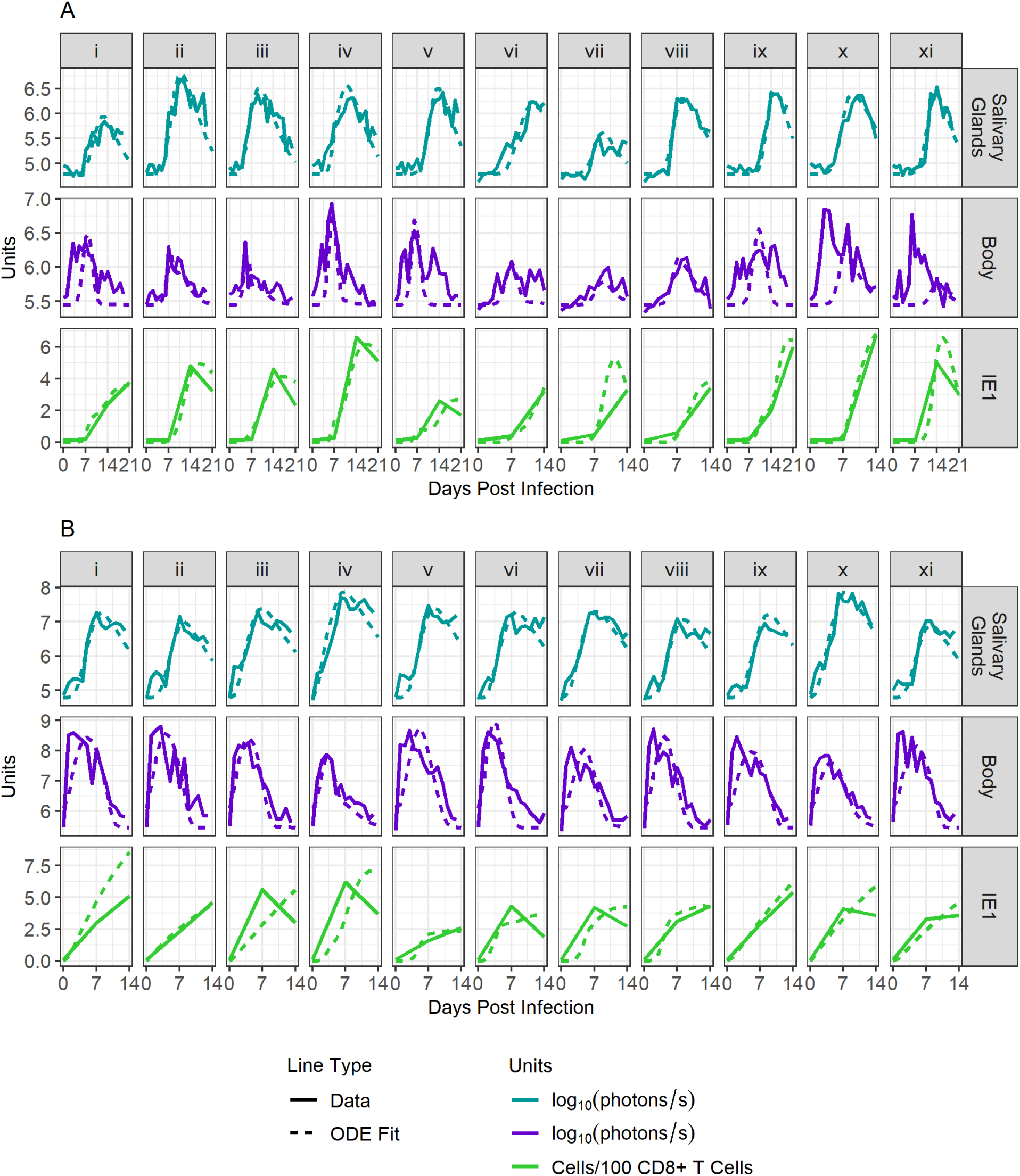
Fits to mice infected IP with K181-luc. Panel A shows model fits for data from mice infected with 100 PFU while panel B shows model fits for data from mice infected with 1,000,000 PFU.

## Notes

### Competing Interest Statement

SG: consulting fees from Moderna, Merck, GSK, Seqirus and Curevo, and research support from Moderna, Merck, GSK, Pfizer, and Altona Diagnostics

